# Nyctinastic thallus movement in the liverwort *Marchantia polymorpha* is regulated by a circadian clock

**DOI:** 10.1101/2020.02.09.940403

**Authors:** Ulf Lagercrantz, Anja Billhardt, Sabine N. Rousku, D. Magnus Eklund

**Affiliations:** Plant Ecology and Evolution, Department of Ecology and Genetics, Evolutionary Biology Centre, Uppsala University, Norbyvägen 18D, SE-75236 Uppsala, Sweden; and the Linnean Centre for Plant Biology in Uppsala.

## Abstract

The circadian clock coordinates an organism’s growth, development and physiology with environmental factors. One illuminating example is the rhythmic growth of hypocotyls and cotyledons in *Arabidopsis thaliana*. Such daily oscillations in leaf position are often referred to as sleep movements or nyctinasty. Here, we report that plantlets of the liverwort *Marchantia polymorpha* show analogous rhythmic movements of thallus lobes, and that the circadian clock controls this rhythm, with auxin a likely meditator. The mechanisms of this circadian clock are partly conserved as compared to angiosperms, with homologs to the core clock genes *PRR*, *RVE* and *TOC1* forming a core transcriptional feedback loop also in *M. polymorpha*.

## INTRODUCTION

Rhythmic movements of plant organs were documented already several centuries BC, but the first known experiments searching for the origin of such rhythms were conducted by the French astronomer de Mairan. Working with a sensitive plant (likely *Mimosa pudica*), he could show that leaves moving in day/night conditions continued to move in constant darkness. During the following centuries, experiments with what Linnaeus later termed “sleep movements” resulted in both the concept of the circadian clock and that of osmotic motors [1,2]. These so called nyctinastic movements often occur in non-growing tissue and are reversible as in several legumes. The reversible movements involve osmotic motors in the pulvinus organ [3], but rhythmic leaf movements can also be growth associated and thus non-reversible. Such rhythms are evident in the movement of leaves in tobacco and cotyledons in *Arabidopsis thaliana* [4,5]. The irreversibility of this process is probably due to deposition of new cell wall material and decreased wall extensibility, but tissue expansion likely results from mechanisms in common with those in pulvinus tissue [6].

Since the introduction of the concept of a circadian or endogenous biological clock great progress has been achieved in understanding the mechanisms behind such internal rhythms. In plants most of this work has been performed in the flowering plant Arabidopsis [7]. A working model of the plant circadian clock comprises a self-sustaining oscillator with an approximately 24-hour rhythm resulting mainly from transcriptional and translational feedback loops [8]. In short, the main components in such models are a set of single MYB domain transcription factors, a family of PSEUDO-RESPONSE REGULATORs (PRRs), and a few plant specific genes with unknown biochemical function. The early morning phased genes *CIRCADIAN CLOCK-ASSOCIATED 1 (CCA1)* and *LATE ELONGATED HYPOCOTYL (LHY)* encode two MYB-like transcription factors that function mainly as repressors of day- and evening-phased genes [9,10,11,12,13]. A second sub-family of related MYB-like transcription factors including *REVEILLE4* (*RVE4*), *RVE6* and *RVE8* has an opposite function, enhancing clock pace through the activation of several core clock genes [14,15].

The family of *PRR* genes comprise five members in Arabidopsis: *PRR1, PRR3, PRR5, PRR7* and *PRR9*. *PRR1* is also known as *TIMING OF CAB EXPRESSION 1 (TOC1)* that together with CCA1 constituted the first conceptual model of the Arabidopsis clock [16]. The expression of *PRR* genes ranges from morning to evening, with *PRR9* peaking in the morning, *PRR5* and *PRR7* around noon, and *PRR3* and *TOC1* around dusk [17]. PRR proteins are in recent models incorporated as transcriptional repressors of *CCA1/LHY* and other *PRR* genes [12]. An additional crucial component of the circadian clock in Arabidopsis is the so-called evening complex (EC), that consists of three proteins: EARLY FLOWERING 3 (ELF3), ELF4, and LUX ARRHYTHMO (LUX) [18,19,20].

When searching for rhythmic growth patterns in the liverwort *M. polymorpha* we discovered that gemmalings (asexually produced plantlets) displayed rhythmic thallus movement. To identify the nature of this movement and the potential involvement of a circadian clock, we studied the function of putative circadian clock genes and their role in controlling the rhythmic movement via the plant hormone auxin.

## RESULTS

### The *M. polymorpha* circadian clock controls nyctinastic thallus movements

In Arabidopsis, growth rates of both hypocotyls and leaves are rhythmic and under the control of a circadian clock [21,22]. In our attempts to detect and measure similar rhythmic growth patterns in liverworts, we noticed that young *M. polymorpha* gemmalings display nyctinastic movements as the lobes of young thalli waves up and down with a 24 h rhythm in conditions of 12 h light and 12 h darkness (neutral days (ND); Figure 1A, B; Supplementary Movie S1). Furthermore, in gemmalings of different accessions these rhythmic movements are maintained in LL (continuous light) conditions with an approximate period of 26.1 – 26.5 h for several days, supporting that they are controlled by a circadian clock (Figure 1C, D, E). One key characteristic of circadian rhythms is temperature compensation, i.e., that the free-running period does not change much with ambient temperature. We thus estimated the free-running period of nyctinastic thallus movement at different temperatures. Consistent with circadian regulation, we found no significant difference in period for temperatures ranging from 18 to 24 °C (Figure 1C).

**Figure 1.**
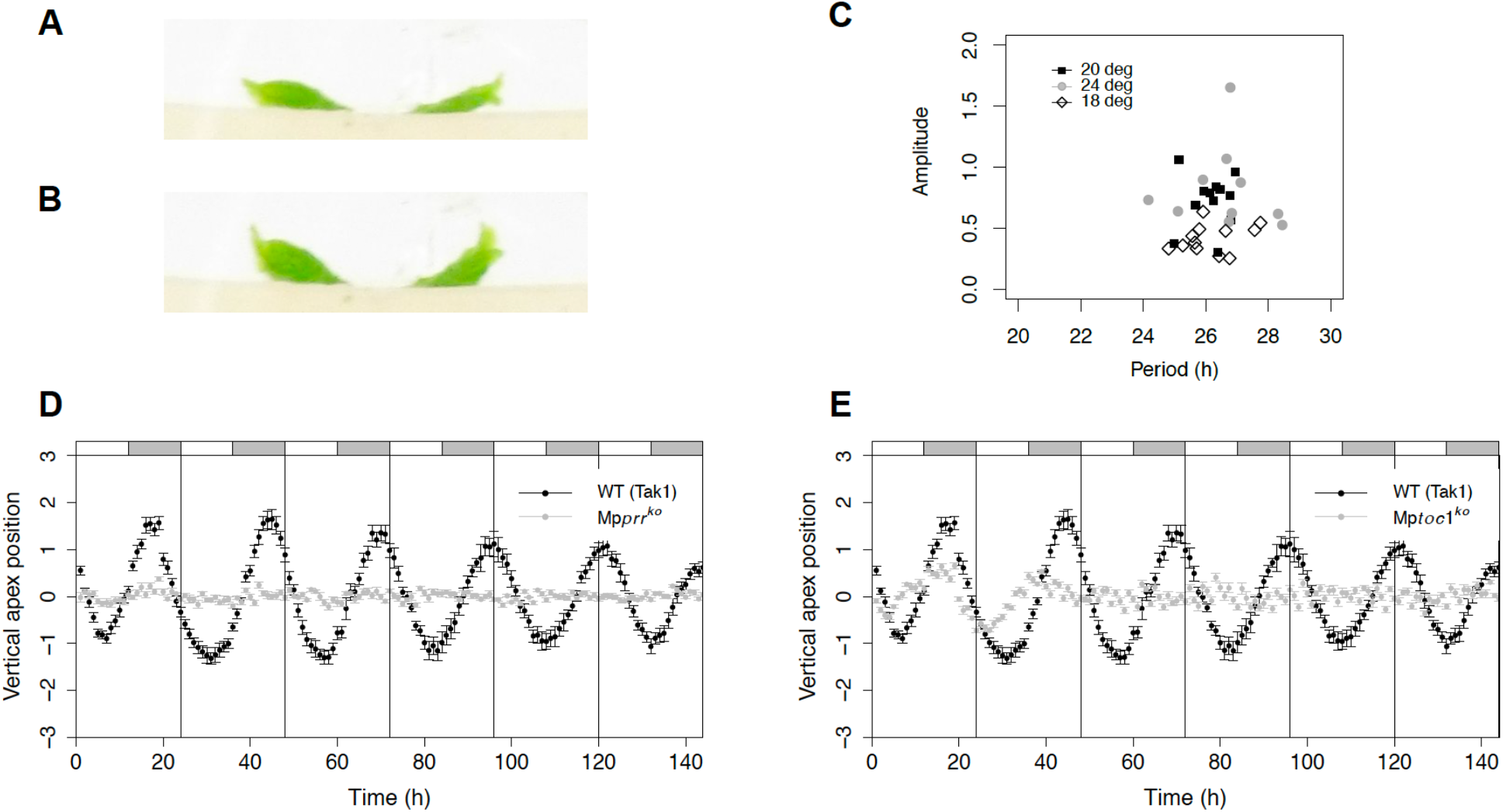
*M. polymorpha* gemmalings display nyctinastic movements of thallus lobes. (A,B) Wild type gemmaling in through (A) and peak (B) positions, respectively. Images were collected at the start and end of subjective day. Gemmalings were grown in a 12h light, 12h dark photoperiod for 5 days, after which light was switched to continuous and imaging was started. (C) Period and amplitude for wild type (Upp10 to 14) at different temperatures. (D, E) Vertical apex position plotted against time for wild type (Tak1, black) and Mp*prr^ko^* (gray) (D), wild type (Tak1, black) and Mp*toc1^ko^* (gray) (E). Apex vertical position data were de-trended using a cubic smoothing spline with 12 degrees of freedom. Data are means ± SE of ten replicate gemmalings.

To further investigate the role of a circadian clock for this movement, we first obtained a more detailed view on the role of Mp*PRR*, Mp*RVE* and Mp*TOC1* as core circadian clock components. Because transcriptional feedback loops are crucial for angiosperm circadian clocks, we examined temporal expression patterns of these genes using qRT-PCR over a 48 h period. As previously reported [23], Mp*PRR* display rhythmic expression in the wild type in LL conditions (Figure 2A). In Mp*toc1^ko^* mutants, expression of Mp*PRR* was continuously high and arrhythmic, as indicated by highly significant effects of both genotype (G) and genotype x time interaction (GxT) terms in ANOVA (*P* < 10^−11^), suggesting that MpTOC1 represses Mp*PRR*. Conversely, expression of Mp*PRR* was low with limited amplitude in Mp*rve^ko^* (*P*-values for both G and GxT terms were < 10^−8^), indicating that MpRVE promote the expression of Mp*PRR* (Figure 2A). Comparing the expression of Mp*TOC1* in wild type, Mp*prr^ko^* and Mp*rve^ko^* similarly suggests MpRVE as an activator also of Mp*TOC1*, and MpPRR a repressor of Mp*TOC1* (Figure 1B; both G (*P* < 10^−4^) and GxT (*P* = 0.026) were significant for Mp*prr^ko^* and G was significant for Mp*rve^ko^* with *P* = 0.017). Furthermore, in Mp*prr^ko^* and Mp*toc1^ko^*, expression of Mp*RVE* remains high and arrhythmic indicating that Mp*RVE* is repressed by both MpPRR and MpTOC1 (Figure 2C; *P*-values for G were less than 10^−9^in both cases, while GxT was marginally significant with *P* = 0.04 and 0.07 for Mp*prr^ko^* and Mp*toc1^ko^*, respectively). These data collectively suggest that Mp*PRR*, Mp*RVE* and Mp*TOC1* are part of a core transcriptional feedback loop of the *M. polymorpha* circadian clock, and that knock-out mutants of these genes can be used to study the role of the circadian clock in the control of growth and development.

**Figure 2.**
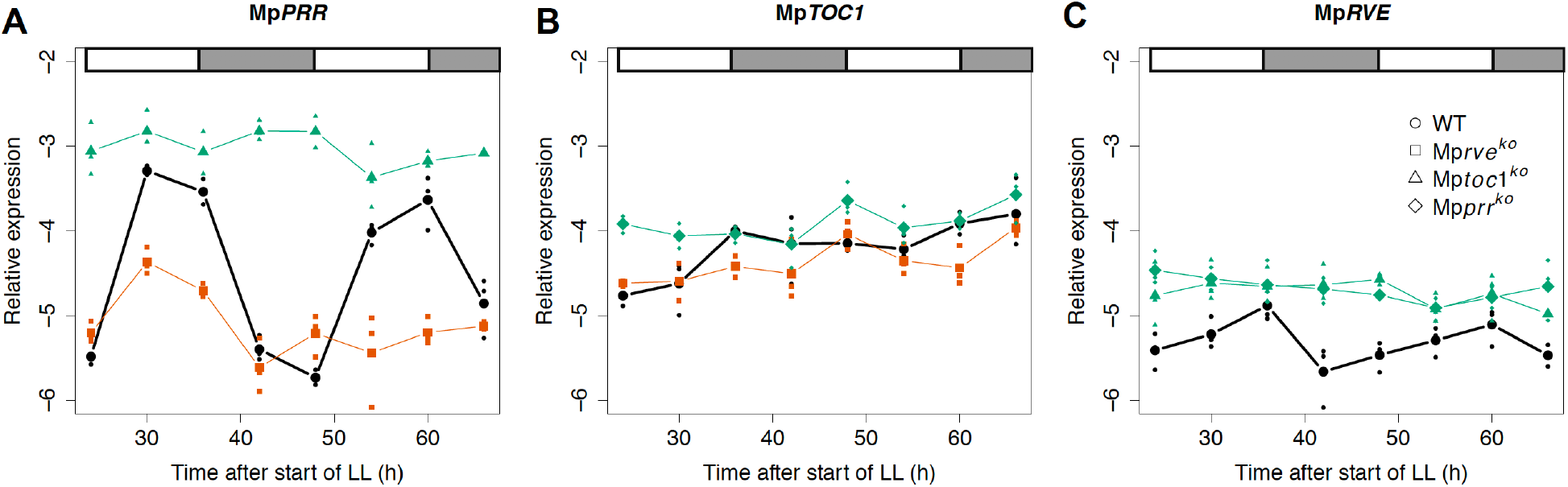
Knockout mutants of Mp*PRR*, Mp*RVE* and Mp*TOC1* affects each other’s expression. qRT-PCRs measuring expression of M. *polymorpha* clock genes during two consecutive days of constant light in wild type (Tak1), Mp*prr^ko^*, Mp*rve^ko^* and Mp*toc1^ko^*. Sampling was conducted every six hours starting 24 hours after switch to LL. Graphs show the expression of (A) Mp*PRR*, (B) Mp*TOC1*, (C) Mp*RVE*. Green and orange lines combined with filled symbols indicate significantly overall higher or lower expression as compared to wild type (bold line), respectively.

We then analyzed rhythms during constant conditions in knockout mutants of Mp*PRR* and Mp*TOC1* that we identified as part of core feedback loops in the *M. polymorpha* circadian clock [23] (Figure 2). In Mp*prr^ko^* and Mp*toc1^ko^* mutants the rhythm is completely lost in LL (Figure 1D, E). These results strongly support that the circadian clock controls “gemmaling waving”, and that estimates of this movement can be used to monitor the *M. polymorpha* circadian clock.

### The circadian clock regulates expression of the auxin biosynthesis gene Mp*TAA*

Most likely, the rhythmic movement is growth related and similar to cotyledon movement in Arabidopsis, suggesting a role for rhythmic auxin production, transport or signaling in its control. Available data suggest that most auxin in liverworts such as *Lunularia cruciata* and *M. polymorpha* is produced in the apical region and transported basipetally through the midrib region producing an auxin gradient [24,25,26,27]. To assay temporal auxin biosynthesis patterns, we analysed gene expression of two genes coding for key enzymes in auxin biosynthesis, Mp*TAA* and Mp*YUC2* [27,28]. In wild-type plants a clear circadian expression pattern was observed for Mp*TAA* (Figure 3A). This pattern was also strongly affected in Mp*toc1^ko^*, Mp*prr^ko^* and Mp*rve^ko^* mutants (Figure 3A). In Mp*prr^ko^* and Mp*toc1^ko^*, expression was reduced and the rhythm dampened, while a higher expression with dampened amplitude was observed for Mp*rve^ko^*. Highly significant effects of genotype (G) was obtained in ANOVA for all cases (*P* < 0.003), while *P*-values for GxT terms were 0.024, 0.009 and 0.053 for Mp*prr^ko^*, Mp*rve^ko^* and Mp*toc1^ko^*, respectively. For Mp*YUC2* expression in the wild type we could not detect a rhythmic pattern in LL, and only Mp*rve^ko^* showed a slightly higher overall expression level with *P* = 0.006 for G term (Figure 3B).

**Figure 3.**
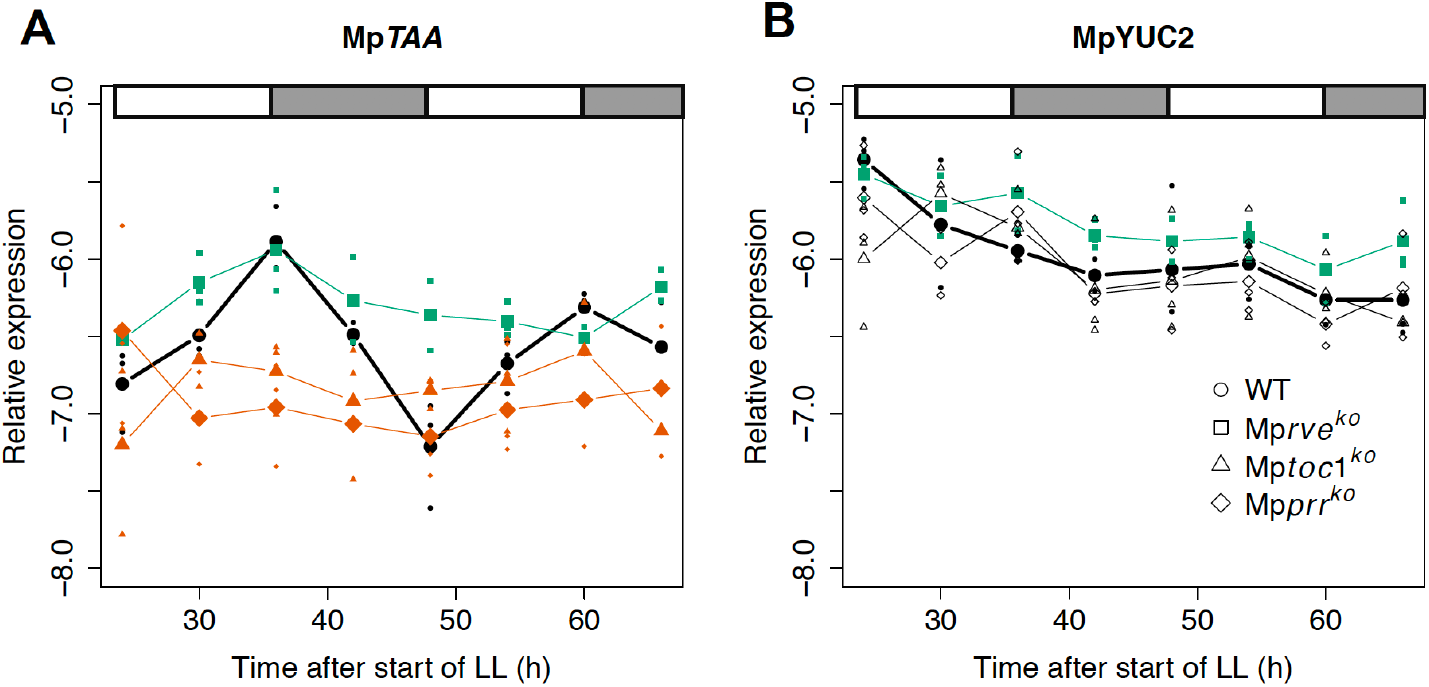
Mp*TAA* display circadian expression that is disrupted in knockout mutants of core clock genes. (A,B) qRT-PCRs measuring expression of *M. polymorpha* auxin biosynthesis genes during two consecutive days of constant light in wild type (Tak1), Mp*prr^ko^*, Mp*rve^ko^* and Mp*toc1^ko^*. Sampling was conducted every six hours starting 24 hours after switch to LL. (A) Mp*TAA* expression. (B) Mp*YUC2* expression. Green and red lines combined with filled symbols indicate significantly overall higher or lower expression as compared to wild type (bold line), respectively.

### Auxin is a likely mediator of circadian control of thallus movements

To further evaluate a role for auxin in the control of rhythmic growth, we conducted nyctinastic movement experiments manipulating auxin levels, distribution and response. For chemical treatments, wild-type gemmae were first grown on standard growth medium for five days to allow for initiation of growth and dorsiventral polarity establishment. Actively growing gemmalings were subsequently transferred to media supplemented with the auxin transport inhibitor 2-[4-(diethylamino)]-2-hydroxybenzoyl benzoic acid [29] (BUM) or low concentrations of indole 3-acetic acid (IAA).

Low doses of IAA (10 and 100 nM) resulted in a reduced angle of growth and thus a more flattened thallus (Figure 4A,C). Rhythmic waving was detectable throughout the experiment on mock and 10 nM IAA, but dampened earlier on 100 nM IAA (Supplementary Movies S2-S4). We cannot exclude that this dampening is due to contact with the solid medium (Supplementary Movie S4). Conversely, an increased growth angle was observed for growth on BUM in a dose-dependent manner (Figure 4B,D). The apparent early dampening of rhythmic waving on BUM could be the result of contact between the two lobes due to the high growth angle.

**Figure 4.**
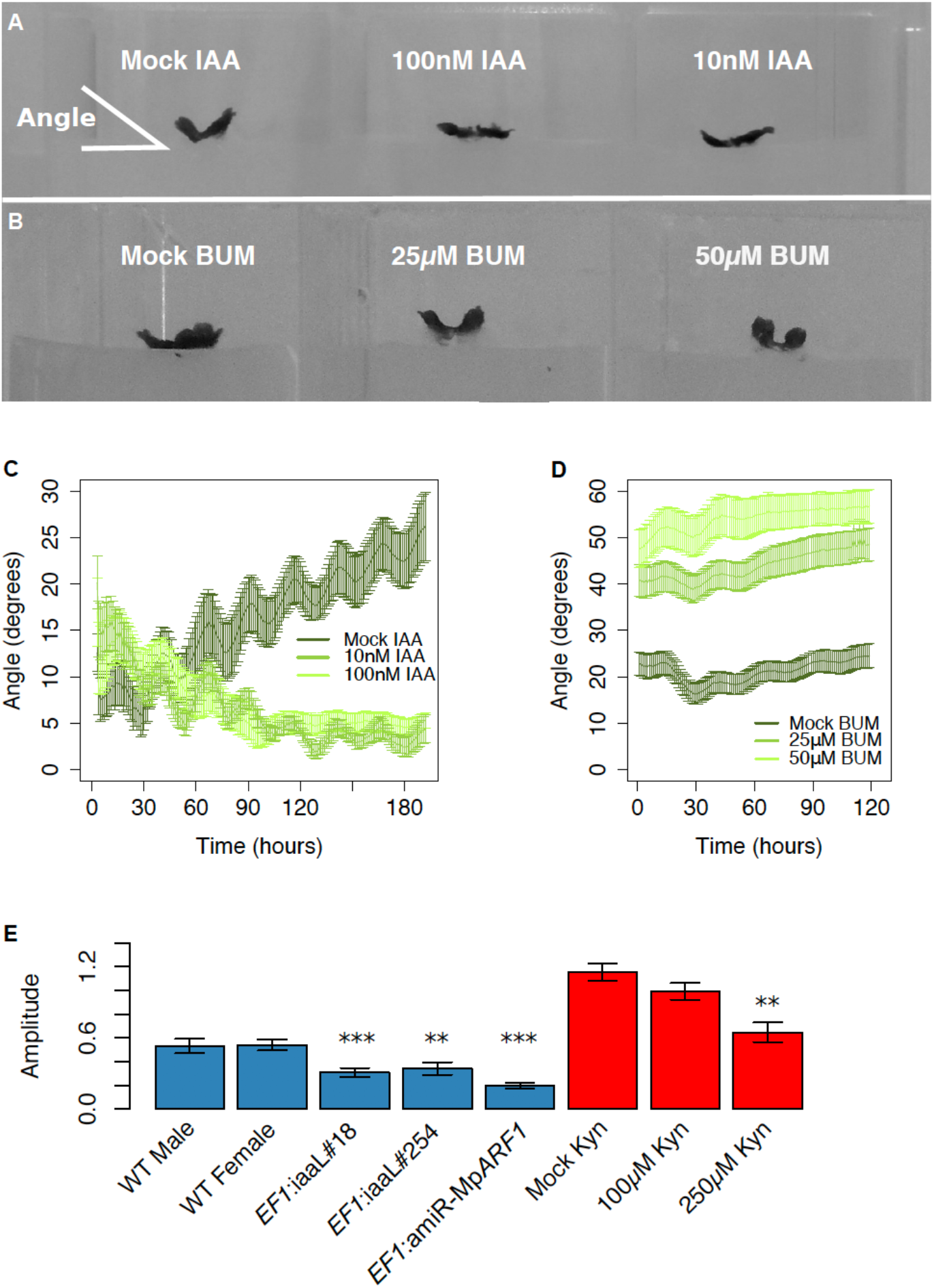
Manipulation of auxin levels, distribution and response affects nyctinastic movements. (A) Gemmalings growing on different concentrations of IAA. (B) Gemmalings growing on different concentrations of BUM. (C) Plot of growth angle over time for gemmalings growing on different concentrations of IAA. (D) Plot of growth angle over time for gemmalings growing on different concentrations of BUM. Angle was measured as indicated in A. (E) Average amplitude of nyctinastic movement for various genotypes and for wild type (Upp5) growing on L-Kyn. ** P < 0.01, *** P < 0.001 for two-tailed t-test against control (WT or mock). WT male and female (E) are of the Australian accession [30]. Graph in E shows means ± SE of three replicates.

To study the effect of reduced auxin levels, we analysed waving in Mp*SHI_pro_:iaaL* plants. These lines express the bacterial auxin-conjugating enzyme IaaL from a promoter mainly active in the apical notch that result in plants with e.g., slow growth and narrow thalli [30]. No difference in period of the rhythmic waving, as compared to wild type, was observed for Mp*SHI_pro_:iaaL* plants, but the analysed line displayed a significantly lower amplitude (Figure 4E). Growth on high concentrations of the TAA inhibitor L-Kynurenine (L-Kyn; 250 μM) gave a similar result; with a significantly reduced amplitude (Figure 4E). Similarly, a reduced amplitude but similar period was observed for *EF1_pro_*:amiR-Mp*ARF1*^Mp*mir160*^ plants [30] (Figure 4E), suggesting that reduced auxin sensitivity also attenuate rhythmic waving. Collectively these results support a role for auxin in rhythmic waving in *M. polymorpha*.

## DISCUSSION

In angiosperms the circadian clock regulates a wide range of processes, including those affecting metabolism, growth, abiotic and biotic stress, and various photoperiodic responses ([31] and references therein). Our previous work suggests an early acquisition of a complex circadian network in plant evolution, but also important differences in the wiring and function of the *M. polymorpha* circadian clock [23].

One unexpected handle of the *M. polymorpha* circadian clock is the rhythmic waving of gemmaling thallus lobes. This movement is most likely related to the non-reversible alternating growth of adaxial and abaxial sides of e.g., cotyledons of Arabidopsis [4]. It has been suggested that this type of nyctinastic movement was ancestral, and that motor cells and pulvini developed later as a means of enhancing leaf movement [32]. If so, our results suggest an early acquisition of nyctinasty already in the first land plants. The cause of this type of alternating growth is poorly known, but may be related to the circadian elongation response of hypocotyls [33]. This rhythmic elongation involves circadian regulated hormone production, transport and/or signaling [34]. Auxin is a good candidate for such a hormone in Arabidopsis and also in *M. polymorpha*.

Our gene expression data suggests that the circadian clock in *M. polymorpha* regulates the first step of auxin biosynthesis, which might contribute to rhythmic auxin levels primarily in the apical region. If auxin levels were to control alternating growth of dorsal and ventral sides we would need to hypothesize a rhythm in dorsal/ventral distribution of the apically produced hormone during the 24-hour cycle. Such a role for auxin in these rhythmic movements is consistent with the results of our manipulations of auxin levels, distribution and response. Application of IAA and BUM had opposite effects on the angle of growth that in turn is directly connected to nyctinastic movements. Assuming that BUM affects ABCB-mediated auxin transport as in Arabidopsis, the increased angle on gemmalings growing on BUM can be interpreted as a result of decreased auxin transport from the ventral to the dorsal side, leading to increased ventral auxin concentration and cell elongation. This hypothesis requires an initial uneven dorsal/ventral auxin distribution, perhaps through a basipetal transport of auxin mainly on the dorsal side. Under this scenario, addition of a significant portion of exogenous auxin (IAA) is expected to result in a more even dorsal/ventral distribution and hence more flat growth (reduced growth angle in our experiments). This is supported by the epinastic growth of gemmalings on high concentrations of auxin or overexpression of the auxin synthesis enzyme MpYUC2 [27,30]. Conversely, L-Kyn-treatment, overexpression of IaaL and the use of amiRNA’s to knock down the expression of auxin biosynthesis genes, resulted in hyponastic growth of gemmalings [27,30]. In our experiments, directly decreasing the amount of active auxin by overexpressing IaaL or affecting IPyA synthesis by adding L-Kyn resulted in lowered amplitude of nyctinastic movement, further supporting the role for auxin in these movements.

Our present results support conservation of the function for the *M. polymorpha* homologs of *PRR*, *TOC1* and *RVE*, each with only one copy in *M. polymorpha* (Mp*PRR*, Mp*RVE* and Mp*TOC1*). Each of them seems to be crucial for maintaining a transcriptional feedback loop in constant light conditions. For Mp*PRR* and Mp*TOC1* we also observed abolished circadian nyctinastic thallus movement, verifying the importance of these genes in generating circadian rhythms in *M. polymorpha*. The stronger effects of mutating these genes in *M. polymorpha*, as compared to Arabidopsis, is likely due to the lack of functionally related paralogs in *M. polymorpha*.

Work on green algae suggest that one homolog of the *PRR/TOC1* clade and one of the *CCA/LHY/RVE* clade constituted the core transcriptional feedback loop in the earliest plants [35,36]. Our work thus supports a continuous use of pairs of *CCA/LHY/RVE* clade genes and *PRR/TOC1* genes at the core of plant circadian clock networks. However, the exact function of these genes within the network seems to have varied over time, partly due to addition of copies of existing genes or new genes to the network, or even deletion of core clock genes.

The only homolog in the whole CCA1/LHY/RVE clade present in *M. polymorpha*, Mp*RVE*, belongs to the LCL sub-clade [23], as do *RVE4*, *RVE6* and *RVE8*. Accordingly, the Mp*RVE* gene does not show an early morning expression, nor acute light induction, which is typical of genes in the *CCA1/LHY* sub-clade [23]. In addition our data support a role for Mp*RVE* as transcriptional activator as opposed to the role of *CCA1/LHY* genes as repressors. Thus, Mp*RVE* seems to have retained a function typical for the *LCL* subfamily [23], despite the loss of the *CCA1/LHY* gene that is absent in all liverworts.

Our identification of a circadian regulated thallus movement provides a practical and easy to use tool for further studies of the evolution of plant circadian clocks, including the effects of frequent gene duplication and circadian gene loss observed during land plant evolution [37,23].

## METHODS

### Plant growth and cultivation

*Marchantia polymorpha* ssp. *ruderalis* Swedish accessions Uppsala (Upp) 1, 5 and 10 to 14, as well as Australian male and female [30], and Takaragaike (Tak)-1 and Tak-2 were grown aseptically on agar solidified Gamborg’s B5 medium [38] (PhytoTechnology Laboratories, Lenexa, KS, USA), pH 5.5. Plants were grown under cool white fluorescent light (50–60 lmol photons m ^2^ s ^1^) in 16 : 8 h, light : dark cycles at 20 °C or as otherwise stated in the text.

### Gene expression analysis

RNA was extracted using an Rneasy Plant Mini Kit (Qiagen). cDNA was synthesized using SuperScript III Reverse Transcriptase (Thermo Fisher) and analysed by qRT-PCR as previously described [23]. Primers are listed in Supplementary Table S1. Mp*EF1α*, Mp*ACT* and Mp*APT3* were used for normalization [39].

For sampling of RNA we used biological replicates – plants of individual transgenic lines or individual wild type lines, and technical replicates – pools of individually grown plants of the same transgenic line or wild type line. We did one cDNA from each RNA sample. In all sampling we used large gemmalings harboring adult tissues, but with no visible gemma cups.

For time series experiments with wild type (Tak1), Mp*prr^ko^*, Mp*rve^ko^* and Mp*toc1^ko^*, three replicate samples of each entity were harvested at six-hour intervals from the second day of LL for two days (in total eight time points). Test of statistically significant expression differences between lines were performed with a linear model in R [40] (aov). The model included time, genotype and their interaction.

### Analyses of nyctinastic thallus movements

25-well square petri dishes (Fisher Scientific) were filled with Gamborg’s B5 medium, after which half of the medium in each well was removed to allow placement and growth of gemmalings. One gemmaling was placed in each well, and plates were placed vertically in a Sanyo growth cabinet (MLR-350) to allow imaging from the side (Supplementary Figure S1). Light was supplied from either cool white fluorescent light, or blue and red LEDs at 20 °C constant temperature, or at the temperatures indicated in the text. Plants where entrained for three to five days in ND (12 : 12 h, light : dark cycles) before exposure to constant conditions and imaging. For auxin related experiments, plants were entrained in ND for one (IAA) or three (BUM) additional days before transfer to LL. Apex position data was extracted from images using ImageJ [41]. Images were converted to binary ones and a rectangular selection automatically following the apex horizontally was used with the command “Analyse Particles” to extract the center of mass in each image. The obtained data on vertical position were de-trended using a cubic smoothing spline with 12 degrees of freedom with the R package smooth.spline.

## ACKNOWLEDGEMENTS

We are grateful for the technical support of Kerstin Jeppsson and Yvonne Meyer-Lucht (EBC, Uppsala University). Tak-1, Tak-2, Mp*prr*^ko^, Mp*toc1*^ko^and Mp*rve^ko^* plants was a gift from Takayuki Kohchi (Kyoto University, Japan). This study was financed by the Swedish Research Council, VR (projects 2014-05220 and 2016-05180 to UL and DME, respectively) and Carl Tryggers Stiftelse för Vetenskaplig Forskning (project CTS17:132 to DME).

## AUTHOR CONTRIBUTIONS

UL and DME designed the research; UL, AB, SNR and DME performed research and analyzed data; UL and DME wrote the paper.

## SUPPLEMENTARY INFORMATION

Supplementary Figure S1. *Marchantia polymorpha* gemmalings growing in square petri dish for waving time-lapse photography.

Supplementary Table S1. Oligonucleotides used in this study.

Supplementary Movie S1. Wild-type *Marchantia polymorpha* gemmaling displaying rhythmic circadian movement of thallus lobes.

Supplementary Movie S2. Wild-type *Marchantia polymorpha* gemmaling growing on media supplemented with mock (control for movies 3 and 4).

Supplementary Movie S3. Wild-type *Marchantia polymorpha* gemmaling growing on media supplemented with 10 nM IAA.

Supplementary Movie S4. Wild-type *Marchantia polymorpha* gemmaling growing on media supplemented with 100 nM IAA.

## DATA AVAILABILITY

The datasets generated during and/or analysed during the current study are available from the corresponding author on reasonable request.

