## Supplementary material for "Nyctinastic thallus movement in the liverwort *Marchantia polymorpha* is regulated by a circadian clock": Supplemenary figure S1

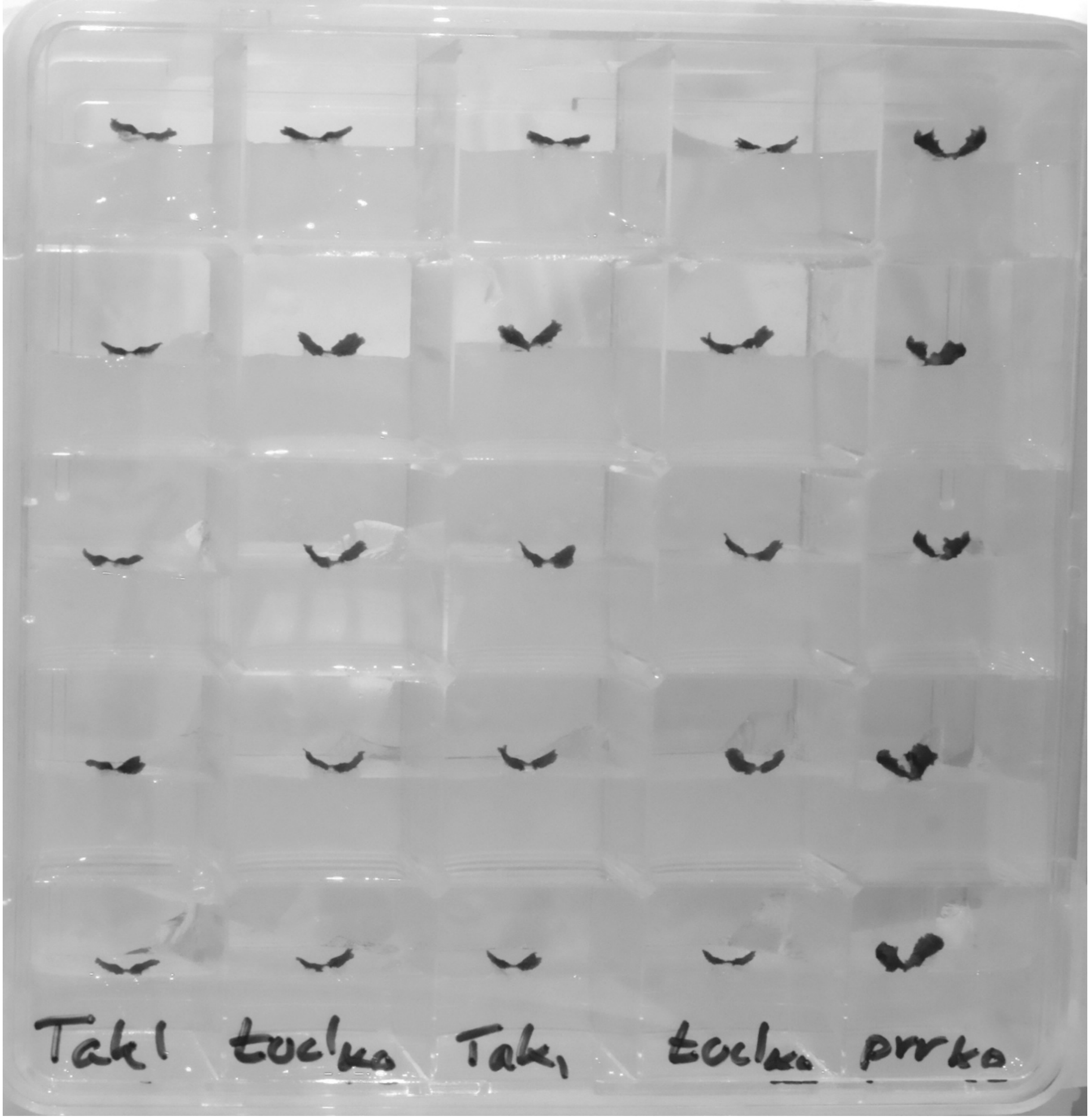


**Supplemental Figure 1** . *Marchantia polymorpha* gemmalings growing in a vertically positioned 25-well square petri dish. Such plates were placed in a Sanyo growth cabinet for imaging of nyctinastic movements.

**Supplemental Table 1.** Oligonucleotides.

| Primer name | Sequence - 5' to 3' | Comments |
|---|---|---|
| ME367 | CGAAAGCCCAAGAAGCTACC | Fwd MpAPT qRT-PCR[1] |
| ME368 | GTACCCCCGGTTGCAATAAG | Rev MpAPT qRT-PCR[1] |
| ME369 | AGGCATCTGGTATCCACGAG | Fwd MpACT qRT-PCR[1] |
| ME370 | ACATGGTCGTTCCTCCAGAC | Rev MpACT qRT-PCR[1] |
| ME402 | CTTGGTTGACTTTGGGCAAT | Fwd MpYUC2 qRT-PCR |
| ME403 | CCGACCTTGTCTTTCAGCTC | Rev MpYUC2 qRT-PCR |
| ME665 | CCGAGATCCTGACCAAGG | Frw MpEF1 qRT-PCR[1] |
| ME666 | GAGGTGGGTACTCAGCGAAG | Rev MpEF1 qRT-PCR[1] |
| ME744 | GGACCAAGTGATTCGCTCTC | Frw MpTAA qRT-PCR |
| ME745 | ACAATGCAGCCTGGAAGAGT | Rev MpTAA qRT-PCR |
| MpPRR F | CAGCAGCTCCTTTGAACAAACA | qRT-PCR[2] |
| MpPRR R | GCCGTGAAGCAGGAAAGAGAAT | qRT-PCR[2] |
| MpRVE F | AAACCTCGGCAAAATCAGGAGT | qRT-PCR[2] |
| MpRVE R | GGCGAGGCAATTTTCAAAGCTG | qRT-PCR[2] |
| MpTOC1 F | CGAAGGAAGAACGACTGAAGCA | qRT-PCR[2] |
| MpTOC1 R | TCTGAGACATTTGACGACGACA | qRT-PCR[2] |

Notes


1. Saint-Marcoux D, Proust H, Dolan L, Langdale JA. 2015. Identification of reference genes for real-time quantitative PCR experiments in the liverwort *Marchantia polymorpha*. *PLOS ONE* 10: e0118678.
2. Linde A-M, Eklund DM, Kubota A, Pederson ERA, Holm K, Gyllenstrand N, Nishihama R, Cronberg N, Muranaka T, Oyama T, Kohchi T, Lagercrantz U. 2017. Early evolution of the land plant circadian clock. *New Phytologist* 216: 576-590.
